# Reproductive success mediates the effects of climate change and grassland management on plant populations dynamics

**DOI:** 10.1101/2023.04.26.538388

**Authors:** Martin Andrzejak, Tiffany M. Knight, Carolin Plos, Lotte Korell

**Author notes:** **Open research statement** The data is currently available at figshare: https://figshare.com/projects/Population_growth_rate_climate_change_and_land_use/157925 and the R code is currently available on GitHub: https://github.com/Martin19910130/Climate_change_and_land_use_population_growth_rate Upon acceptance the data will be available at figshare: https://figshare.com/projects/Population_growth_rate_climate_change_and_land_use/157925 and the R code on GitHub: https://github.com/Martin19910130/Climate_change_and_land_use_population_growth_rate.

## Abstract

Climate change is one of the largest threats to grassland plant species, which can be modified by land management. Although climate change and land management can separately and interactively influence plant demography this has been rarely considered within one experimental set-up. We used a large-scale experiment to quantify the effects of grassland management, climate change and their joint effect on the demography and population growth rate of 11 native plant species. We parameterized integral projection models with four years of demographic data to project the population growth rate. We hypothesized, plants would perform better in ambient than in the future climate treatment that creates hotter and drier summer conditions and that plant performance in grazing vs. mowing would vary across species and depend on their traits. Due to extreme drought events, over half of our study species went quasi extinct, which highlights how extreme climate events can influence long term experimental results. Of the persistent species, only one supported our expectations, and the rest either had higher population growth rates in the future climate treatment or showed no significant difference in population growth between climate treatments. Species with shorter flowering durations performed better in the mowing treatment while those with longer flowering durations performed better in the grazing treatment. The population growth rates of these species were sensitive to changes in vital rates related to reproduction and recruitment. Depending on the species we found effects of land management and climate change on population growth rates but we did not find strong support for interactive effects among both factors. Experiments combined with measuring plant demographic responses provide a way to isolate the effects of different drivers on the long-term persistence of species, and to identify the demographic vital rates that are critical to manage in the future. Our study suggests that it will become increasingly difficult to maintain species with preferences for moister soil conditions, that traits such as flowering duration might predict responses to management, and that vital rates such as reproduction and recruitment are disproportionately important.

## Introduction

Grasslands cover approximately 40% of the Earth’s surface (Bardgett et al., 2021) and provide important ecosystem services to society, such as food for livestock and humanity, pollination and biodiversity (Biurrun et al., 2021; Feurdean et al., 2018). In Central Europe, semi-natural grasslands were historically managed using low-intensity (i.e. extensive) mowing or grazing, which prevents their succession into forests (Poschlod et al., 1998; Tälle et al., 2015). These different types of management affect which types of plants can persist. For example, grazing favors plants with traits related to higher grazing resistance (e.g. hairiness) or grazing tolerance (e.g. resprouting capacity) (Bernhardt-Römermann et al., 2011; Evju et al., 2009; Kahmen et al., 2002). Anthropogenic climate change may modify the effects that management has on the demography and persistence of plant species (Bardgett et al., 2021; Kariyeva et al., 2012; Redlich et al., 2022). For example, individuals that currently avoid loss of reproduction from mowing by flowering before or after mowing events might shift in their phenology in the future due to climate change, which could lead to mismatches between flowering timing and mowing events that harm plant reproductive success (Cleland et al., 2006; Gordo & Sanz, 2010).

In central Europe, the climate is projected to become warmer, wetter in spring and fall, and drier in summer in the future (Döscher et al., 2002; Jacob & Podzun, 1997; Rockel et al., 2008; Wagner et al., 2013). Drought-tolerant species should be best able to survive the drier summers expected in the future (Belovsky & Slade, 2021). Species that flower earlier and across a longer time period might be favored in the future, because they can take best advantage of the wet spring conditions and because the long flowering duration might buffer their populations from complete reproductive failure during periods of extreme weather. This leads to the expectation that drought-tolerant plants with an early flowering start combined with a long flowering period should be best able to avoid losing reproductive success under future climate conditions in the presence of grazing and mowing land management types.

In order to understand population dynamics of plant species, demographic studies quantify vital rates such as survival, growth and reproductive output across the entire life cycle of the plant and project long-term population growth rate using structured population models (Caswell, 2000; Crone et al., 2011; Easterling et al., 2000). A common approach to analyzing plant demographic data are Integral Projection models (IPMs), which consider a relationship between continuous plant size and vital rates and can also incorporate discrete development stages (Childs et al., 2003; Jacquemyn et al., 2010; Yule et al., 2013). Retrospective analyses of IPMs, such as life table response experiments, decompose the contribution of individual vital rates to the changes in population growth rate between treatments. Vital rates that are more responsible for the change are those that, either change dramatically between the treatments and/or those that the population growth rate is sensitive to changes in (Rees & Ellner, 2009).

Many demographic studies have been conducted to assess how the demography of species responds to climate (Ehrlen, 2019; Ehrlén et al., 2016; Ehrlén & Morris, 2015; Lemmer et al., 2021; Morris et al., 2020). Most studies focus on natural variation across space and time in climate conditions and measure plant demographic responses (Compagnoni et al., 2021). Such observational studies are limited in their ability to assess cause and effect relationships, as there are often other important factors that co-vary with climate. A few studies have experimentally assessed how climate affects plant demography (Compagnoni & Adler, 2014; Lemmer et al., 2021; Lyu & Alexander, 2022; Töpper et al., 2018; Williams et al., 2007). However, most climate change experiments consider extreme manipulations of temperature and precipitation, making it hard to predict how plants might respond under realistic future climate change scenarios (Korell et al., 2020).

We have a unique opportunity to study how a realistic future climate scenario as well as two different grassland management types (extensive grazing vs. mowing) influence the demography and population dynamics of plants using the Global Change Experimental Facility (GCEF) in Germany. Our study builds on the research of Lemmer and colleagues (Lemmer et al., 2021), which discovered that land management types and climate change interactively affected the population growth rate of the drought-tolerant grass, *Bromus erectus* L., and that this interaction was primarily due to higher seedling recruitment of plants under the future climate and mowing management treatment combination. Lemmer et al. (2020) measured the demography across a short time period (one transition, two years) and concentrated on a single species. In this study, we expand the scope to understand the demography and population responses to climate and land management types across multiple transitions along a longer time period and multiple plant species. Demographic data was collected over the course of five years (2018 – 2022) for eleven (Table 1) grassland plant species that represent a range of different plant statures (tall vs. small), life forms (grasses, herbs) and life histories (biennials vs perennials). In general, we expected i) the effects of land management types will vary across species and depend on their traits, ii) plants show higher population growth rates under ambient compared to future climate conditions and iii) the effects of climate treatments and land management types on the population growth rate will be interactive.

**Table 1.**
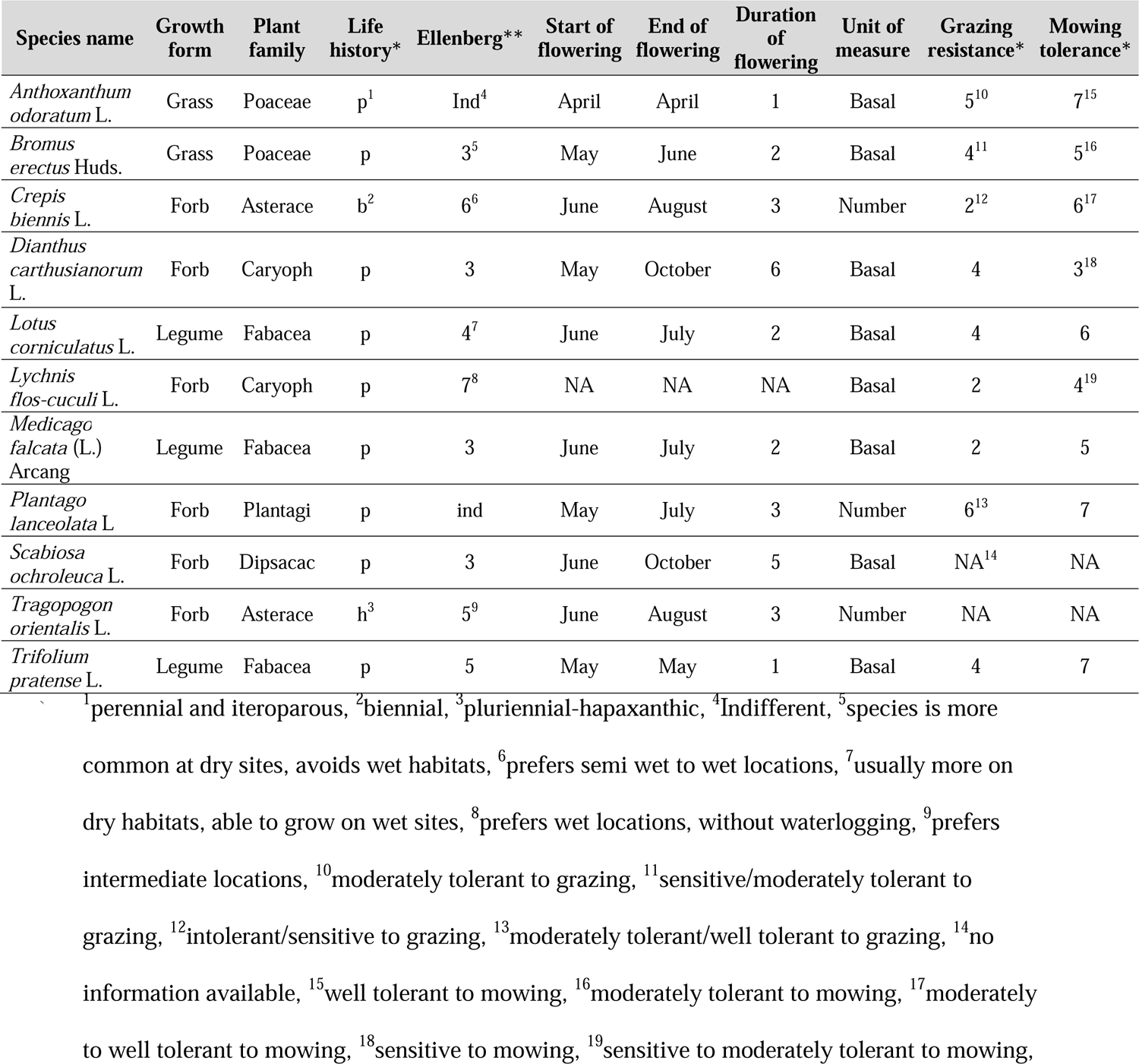
Description and ecology of each plant species in the study as extracted from the following sources: https://wiki.ufz.de/biolflor/index.jsp* and Ellenberg et al., 1991**.

| Species name | Growth form | Plant family | Life history* | Ellenberg** | Start of flowering | End of flowering | Duration of flowering | Unit of measure | Grazing resistance* | Mowing tolerance* |
| --- | --- | --- | --- | --- | --- | --- | --- | --- | --- | --- |
| *Anthoxanthum odoratum* L. | Grass | Poaceae | p<sup>1</sup> | Ind<sup>4</sup> | April | April | 1 | Basal | 5<sup>10</sup> | 7<sup>15</sup> |
| *Bromus erectus* Huds. | Grass | Poaceae | p | 3<sup>5</sup> | May | June | 2 | Basal | 4<sup>11</sup> | 5<sup>16</sup> |
| *Crepis biennis* L. | Forb | Asterace | b<sup>2</sup> | 6<sup>6</sup> | June | August | 3 | Number | 2<sup>12</sup> | 6<sup>17</sup> |
| *Dianthus carthusianorum* L. | Forb | Caryoph | p | 3 | May | October | 6 | Basal | 4 | 3<sup>18</sup> |
| *Lotus corniculatus* L. | Legume | Fabacea | p | 4<sup>7</sup> | June | July | 2 | Basal | 4 | 6 |
| *Lychnis flos-cuculi* L. | Forb | Caryoph | p | 7<sup>8</sup> | NA | NA | NA | Basal | 2 | 4<sup>19</sup> |
| *Medicago falcata* (L.) Arcang | Legume | Fabacea | p | 3 | June | July | 2 | Basal | 2 | 5 |
| *Plantago lanceolata* L. | Forb | Plantagi | p | ind | May | July | 3 | Number | 6<sup>13</sup> | 7 |
| *Scabiosa ochroleuca* L. | Forb | Dipsacac | p | 3 | June | October | 5 | Basal | NA<sup>14</sup> | NA |
| *Tragopogon orientalis* L. | Forb | Asterace | h<sup>3</sup> | 5<sup>9</sup> | June | August | 3 | Number | NA | NA |
| *Trifolium pratense* L. | Legume | Fabacea | p | 5 | May | May | 1 | Basal | 4 | 7 |

<sup>1</sup>perennial and iteroparous, <sup>2</sup>biennial, <sup>3</sup>pluriennial-hapaxanthic, <sup>4</sup>Indifferent, <sup>5</sup>species is more common at dry sites, avoids wet habitats, <sup>6</sup>prefers semi wet to wet locations, <sup>7</sup>usually more on dry habitats, able to grow on wet sites, <sup>8</sup>prefers wet locations, without waterlogging, <sup>9</sup>prefers intermediate locations, <sup>10</sup>moderately tolerant to grazing, <sup>11</sup>sensitive/moderately tolerant to grazing, <sup>12</sup>intolerant/sensitive to grazing, <sup>13</sup>moderately tolerant/well tolerant to grazing, <sup>14</sup>no information available, <sup>15</sup>well tolerant to mowing, <sup>16</sup>moderately tolerant to mowing, <sup>17</sup>moderately to well tolerant to mowing, <sup>18</sup>sensitive to mowing, <sup>19</sup>sensitive to moderately tolerant to mowing,

**Table 1.** All vital rates and parameters that where used and extracted to calculate the integral projection model (IPM)

| Vital rates | Abbreviation | Life-cycle stage | Parameters | Distribution | LTRE stage |
| --- | --- | --- | --- | --- | --- |
| Plant survival | S | Survival | Intercept (Int), Slope | Bernoulli | Survival |
| Plant growth | G | Growth | Intercept (Int), Slope, SD | Normal | Growth |
| Reproduction probability | P | Reproduction | Intercept (Int), Slope | Bernoulli | Reproduction |
| Number of seeds per reproductive plant | F | Reproduction | Intercept (Int), Slope | Poisson | Reproduction |
| Fall recruitment | $\theta_f$ | Recruitment | Mean | | Recruitment |
| Spring recruitment | $\theta_s$ | Recruitment | Mean | | Recruitment |
| Seedling establishment | B | Establishment | Mean |  | Establishment |
| Size distribution of new plants | $\eta$ | Establishment | Mean, SD | | Establishment |

## Methods

### Study site

We conducted our research in the Global Change Experimental Facility (GCEF), which was established in 2013. The experiment is part of the research station Bad Lauchstädt (51°22060 N, 11°50060 E, 118 m a.s.l.). It is designed to simulate a realistic future climate scenario combined with the manipulation of different land management types, including extensively managed grasslands. The future climate treatment is based on a mean of 12 regional climate models for the region (Schädler et al., 2019).

The GCEF consists of 10 main units, five with ambient climate and five with future climate conditions. Precipitation in the future climate treatment is manipulated seasonally: increases in precipitation by +10 % compared to the ambient climate in spring and fall and reduced precipitation by −20 % compared to the ambient climate in summer (for more details see Schädler et al. 2019). The minimum temperature in the future climate treatment is increased by about +1.5°C by passive nighttime warming (Schädler et al. 2019).

Each main unit is divided into five subunits that are 16m x 24m in size, each representing a different land management. Each subunit has 14m x 19.5m available for measurements and the remainder is left as an edge area, where no experiments or measurements can be undertaken (Appendix S1, Figure S1). Our study was established in two of the five land managements, the extensive meadow (mown one or two times per year) and the extensive pasture (grazed by 20 sheep for 24 hours twice to three times per year). These two land management types were initially sown in 2013 with 56 plant species native to Germany (Schädler et al., 2019). This design leads to the following treatment combinations in our experiment: (1) ambient grazing, (2) ambient mowing, (3) future grazing and (4) future mowing.

### Study species

In 2018, we chose eleven different species (Table 1) for our demographic study based on the criteria that they were abundant enough for demographic data collection (minimum number of individuals in 2018 was 50 per treatment combination), and that they were species that are typically associated with mesic or dry grasslands. The species covered a range of growth forms, plant families and life histories (biennial vs. perennial). Every plant species in this study is native to Germany and was sown into the GCEF experiment when it was established.

### Demographic data collection

In April 2018, we established one transect on all our study subunits resulting in 20 transects (five replicates per treatment combination). Our goal was to find a minimum of 10 individuals per transect, to reach a minimum of 50 individuals per treatment combination for every species. Along each transect, 50 x 50 cm subplots were established. We assigned three of the subplots to a fixed positions along each transect (0– 0.50 m; 3 – 3,50 m; 6-6.50 m). The location of other subplots were chosen along the transect at strategic locations to capture individuals of species that were needed to achieve our target of 50 individuals per species and transect. Following this procedure, most transects had eight subplots. This leads to a different overall subplot number for each species, with rarer species having more subplots than common species. Each subplot was marked with a land marking (pipe with red plastic top) in the bottom left and a nail with a white plastic ring the top right corner.

In April 2018, we measured the size of each individual target species in each subplot using either basal area or number of leaves (Table 1). Basal area (cm^2^) measured as the product of the horizontal width and length at the base of each individual. Length was measured as the longest lateral dimension of each plant, and width was measured perpendicular to the length. For rosette species (e.g. *Plantago lanceolata, Tragopogon orientalis*), we used the number of leaves as proxy for their size. We recorded the location of each individual within the 50 x 50 cm subplot, using an XY-coordinate system so that individuals could be followed over the years. In rare cases if we spotted individuals of our rarer species just outside of our subplot, we would measure those individuals and track their position relative to the subplot (one of the coordinates would exceed 50 cm).

From 2019 until 2022, we went to the field five times a year. In April, we re-located individuals from the previous year and measured their size, identified new individuals and measured their size, and counted the number of new seedlings (for details on how we identified seedlings see Appendix S3, Table S2). Depending on the phenology of the species and the management events (see Table 1, Table S1), we counted flowers and seed heads of all reproductive individuals in each transect. Further we collected two seed heads per subunit and species from individuals that were outside of our transects but within the same subunits. Number of seeds in each seed head were counted in the lab. In late autumn (October/November), we counted the number of seedlings in our subplots, as seedlings of these species can germinate in spring or in fall.

From 2019 onwards, we decided to mark individuals of the very abundant species *Bromus erectus* with ID’s as that helped separating and finding individuals. To mark the individuals, we used rings made of wire with a tag with a running number on it and put it around the base of the individual. We left enough space for the individual to grow. Special care was taken to not harm any of the vegetation during that process.

The multiple droughts Germany experienced during several of the years of this study prevented more frequent mowing and grazing than would otherwise be incorporated into the treatment plan (Schädler et al., 2019). During the time of our experiment, the extensive meadows were mown once per year in early summer time from 2018 – 2020 and two times (early and late summer) in 2021. The extensively grazed subplots were grazed twice per year in spring and early summer from 2018-2020, and three times in 2021 (spring, early and late summer). For a detailed look at the management timing, see Appendix S2, Table S1.

For information on traits for our species, we used Ellenberg values (Ellenberg et al., 1991), the Bioflor database (https://wiki.ufz.de/biolflor/index.jsp) and our own measurements of phenology. For drought tolerance, we used categorical Ellenberg values (Ellenberg et al., 1991, Table 1). For grazing and mowing resistance, we used categorical values from the Bioflor database (Table 1). In 2020, we collected phenology data on our target species in the GCEF between April and December within the central 3 x 3 m vegetation survey plot of each subunit where no experiments or other disturbances (besides the management events) occur. This provides information on the month in which flowering began and the month it ended, taking all individuals inside the central vegetation plot into account.

### Quasi extinction

Some of the species declined so significantly over the course of our demographic study, that we considered them quasi extinct and no longer continued to collect demographic data. We defined quasi extinction from a treatment combination as fewer than 25 individuals total and fewer than 10 flowering individuals, thus those species treated the species as quasi extinct.

### Life cycle stages and vital rates

For all species, we modeled a year-to-year life cycle (April to April) with two stage classes. A discrete class for seedlings and a continuous class for plants (Figure 1). Most of our species do not form long lasting seed banks, and thus we do not have a seed bank class. *Plantago lanceolata* does have a long-lived seed bank (Chen, 2018), but most of the seeds of this species germinate immediately and therefore demographic studies on this species do not typically incorporate the seed bank stage (e.g., www.plantpopnet.com).

**Figure 1.**
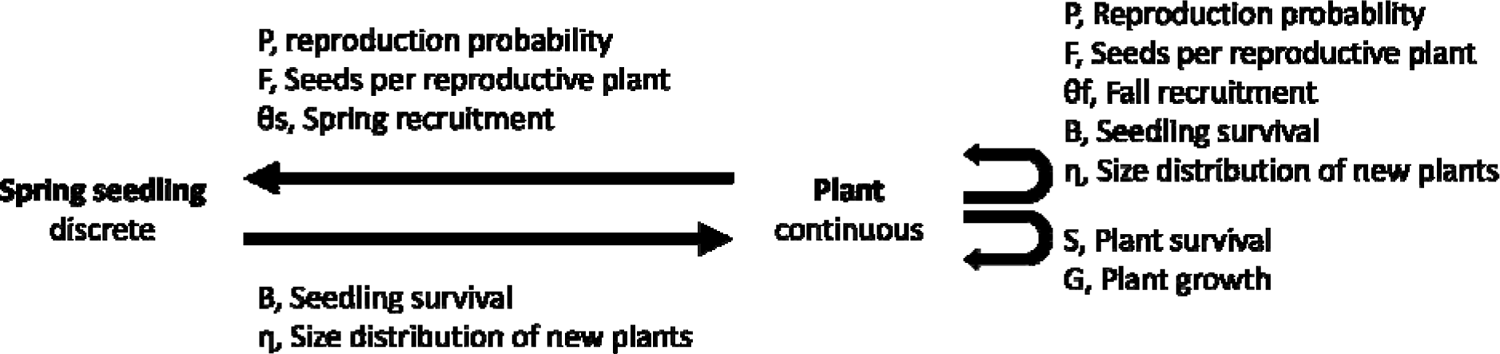
Life cycle diagram of the study species. The diagram shows every vital rate incorporated in the integral projection model (IPM) and the abbreviation used.

We modeled each continuous state of the IPM as a function of the natural logarithm of the size of an individual. We used generalized linear models to fit all continuous variables. Survival is a function of whether or not an individual (i) was still alive at t1, dependent on its log transformed size (z) at t0 (Eq. 1), modeled as a Bernoulli process with probability of survival S_t1_ (Eq. 2).

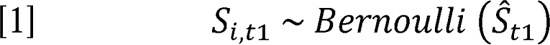

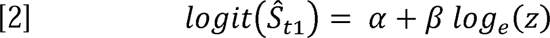

In this function α is the intercept and β the slope of the curve.

Growth of plants G_i, t1_ is described as the normally distributed change in z and log transformed size at t1 (z’) of a surviving individual plant (i) (Eq. 3). It was modeled as a linear function of z’ of surviving individuals depending on z, where the intercept is defined as α^G^, the slope as β^G^ (Eq. 4) and the standard deviation as σ_G_ (Eq. 3).

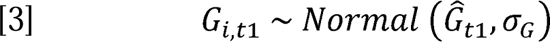

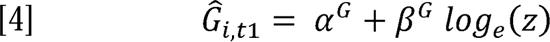

Similar to survival, we modeled reproduction probability (P_i, t0_) as Bernoulli process (Eq. 5). Here we tested how the probability an individual i reproduces at time t0 depends on z. The intercept is defined as α^P^ and the slope as β^P^ using a logit link function (Eq. 6)

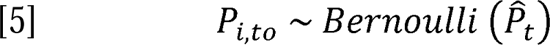

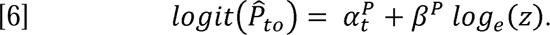

For every reproductive plant, we calculated the number of seeds it produced (Eq. 7). The number of seeds produced by an individual I at t0 was a function of z, modeled as a Poisson distribution. The intercept was defined as α^F^ and slope β^F^ (Eq. 8)

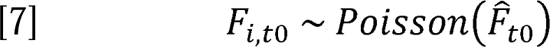

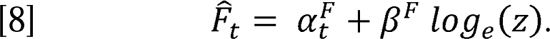

Recruitment, the emerged seedlings per total number of seeds, was separated into fall - (θ*_f, j, t0_*) and spring recruitment (θ_s,_ *_j, t0_*). To calculate recruitment, we summed the total number of seeds per subplot (S_j,t_, Eq. 9). Where S_i,t1_ is the total number of seeds produced by one individual (n) in a subplot at time t1.

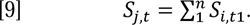

We then divided the result of Eq. 9 by the number of seedlings found in that subplot in fall t0 (*Rf_j, to_*, Eq. 10) and spring t1 (Rs_j,t1_, Eq. 11).

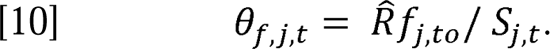

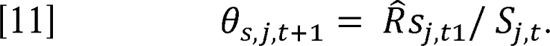

Establishment (B_j_) is the proportion of seedlings that survive to become plants from t0 to t1 in each subplot j. We calculated the sum of seedlings in spring and fall t0 (Rsum_j, t0_) of a subplot and divided that number by the number of new individuals at t1 (Ni_j, t1_) in the same subplot.

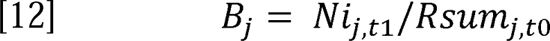

We calculated the log size distribution of new plants η which is defined as the normally distributed size of individuals that entered the continuous plant stage in year t1 (Eq. 12). We used the mean (loge(ηrtl)) and standard deviation (σ_η_) of this size in our final parametrization of the IPM.

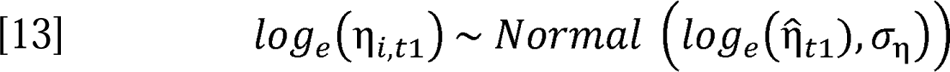

### Effects of treatments on vital rates

For each vital rate (survival, growth, reproduction probability, number of seeds, recruitment, establishment, size distribution of new plants), we generated three independent pipelines. The first pipeline differentiated between the climate treatment and thus we calculated the parameters for ambient and future climate, respectively (Table 2). The second pipeline differentiated between land management types (mowing and grazing). In the third pipeline, we differentiated between all four treatment combinations to also account for possible interactions among treatments. In each of those models, we dropped year, to make the models more robust (but see Appendix S5 for the “yearly” models).

### Integral projection model (IPM)

For each species and pipeline, we created an IPM to calculate the influence of climate and management on the long-term population growth (λ). In every IPM, we calculate all possible transitions from z to z’ at. The change in number of plants from one year to the other is described by:

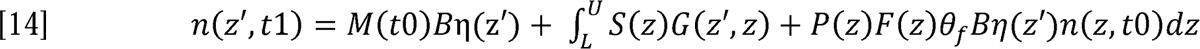

Where the vector n(z’, t1) is the number of plants at size z’ at time t1. In this function, the first term represents the recruitment of spring seedlings entering the adult plant class. This is based on the number of spring seedlings at t0, M(t0), the seedling establishment, B, and the size distribution of new plants η(z’). The second kernel (or surface) describes the growth and survival of plants from t0, n(z, t0) to t1, n(z’, t1). It is defined as an integral with in the upper limit U, which is the biggest size observed and the lowest size L, observed for the species in our study. The integrals were evaluated across 200 equally spaced size bins using the midpoint rules as a 200 x 200 matrix. The recruitment of spring seedlings from one year to the next is described by:

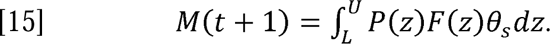

As we did not distinguish the new plants we see in April as those that were seedlings from fall vs. spring, we use the same establishment estimate for seedlings (B) twice in the model.

### Effects of treatments on population growth rate

To determine the effect of the climate treatment on λ, we calculated λ for the ambient and future treatments and then calculated an effect size by subtracting λ ambient from λ future. Positive values show a specieś preference for ambient climate conditions whereas negative values show a specieś preference for future climate conditions. Likewise, for land management types, we subtracted λ grazed from λ mowing to calculate an effect size. Here, positive values show a species preference for the grazing treatment whereas negative values show a species favors the mowing treatment.

To test whether traits influenced the effect size of land management types on λ, we ran simple linear models using the effect size as response variable and the following traits as explanatory variable: flowering start month, duration of flowering, grazing resistance and mowing resistance. Because flowering time, start and end could differ between treatments due to the effect of the land management type we decided to not calculate a mean across those.

To quantify uncertainty in estimates of λ, we bootstrapped our data a thousand times and calculated the λ each time. To test for significant differences in λ between treatments and treatment combinations, we used permutation (randomization) tests (N = 1000 permutations). For each treatment combination, we calculated sensitivities and elasticities of λ to kernel element perturbations.

### Life table response experiment (LTRE)

In order to decompose the influences of vital rates on the difference in λ, across treatment combinations, we conducted Life Table Response Experiments (LTRE) for each pairwise treatment combination. The difference in λ, Δλ, is:

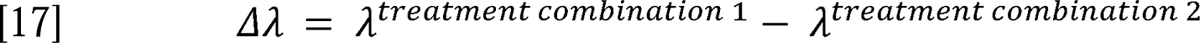

The contribution of each vital rate to Δλ, δ^∼^ ^αi^, is:

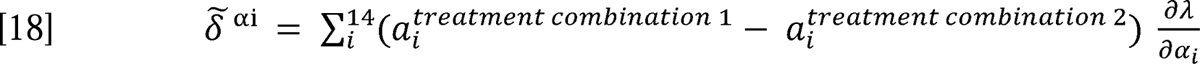

Here α_i_ is one of the fourteen vital rates that are included in the IPMs, while the sensitivity of λ to each vital rate is defined as 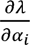. Vital rates will have a strong contribution to the observed α_i_ difference in λ between treatment combinations if the vital rates change strongly in magnitude between the two treatment combinations and/or if λ is sensitive to changes in that vital rate. To facilitate the understandability, we combined the vital rates with respect to six demographic processes: survival, growth, reproduction, recruitment, establishment (Table 2). After this we scaled o^∼^ ^αi^ to 1 (to display proportional influence).

### Analysis

All analysis were carried out in R (R Core Team, 2022). Figures were made using the ggplot2 package (Wickham, 2016). The integral projection model was run and calculated using the ipmr packages (Levin et al., 2021). Other packages used were: openxlsx (Schauberger & Walker, 2021) to load the xlsx files into R, dplyr (Wickham et al., 2022) for structuring data and piping and the package parallel (R Core Team, 2022) to decrease the running time of R scripts.

## Results

### Quasi extinction

The following six species went quasi extinct over the course of the experiment: *Anthoxanthum odoratum, Crepis biennis, Lotus corniculatus, Lychnis flos-cuculi* and *Trifolium pratense*. While there is too small of a sample size (N=11 species) for statistical analysis, it is notable that the species that went quasi extinct were also the ones that were less drought tolerant based on their Ellenberg values (Table 1) and that the first few years of our study were extraordinarily dry for our region (Boergens et al., 2020).

### Management effects on population growth rate

*Tragopogon orientalis* had significantly higher λ in mowing compared to grazing management (permutation test p = 0.042, Figure 2e), whereas all other species did not show a significant main effect of management on λ (Figure 2a-d).

**Figure 2.**
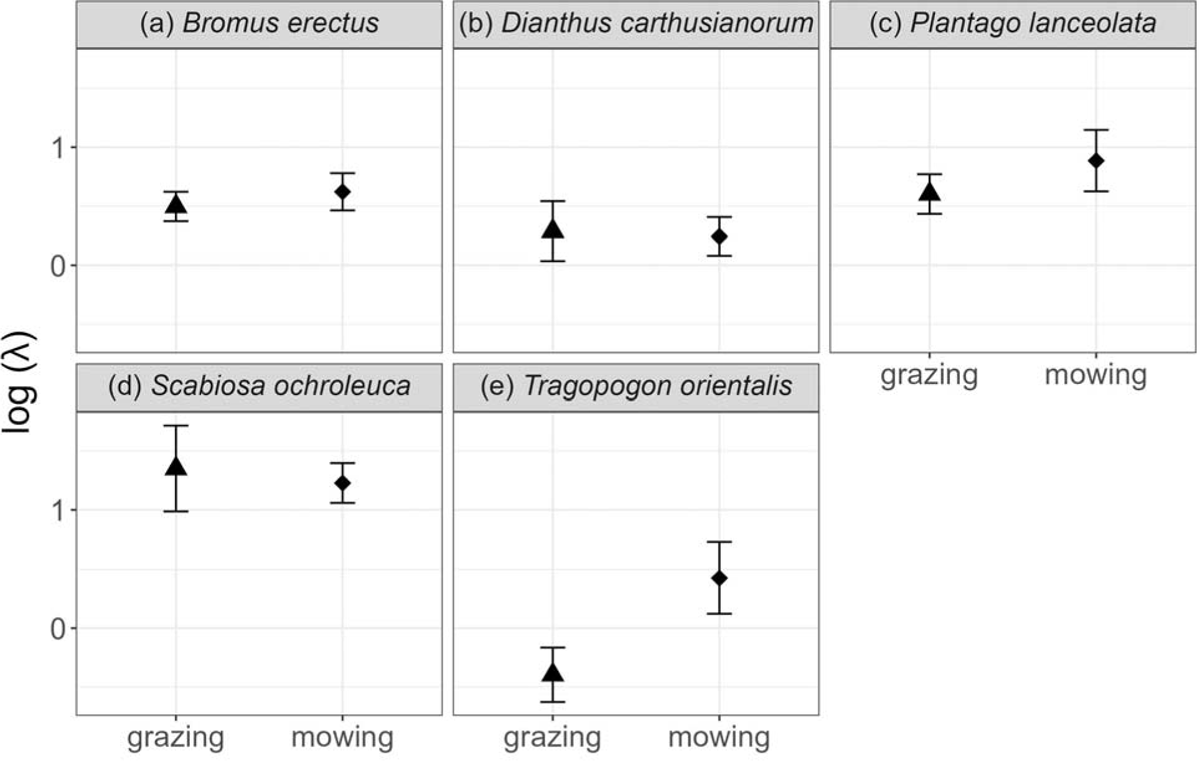
Log population growth rate (λ) for each of the five study species (a-e) considering the two different land management types (grazing and mowing). Displayed is the mean ± one standard deviation.

Traits related to grazing resistance and mowing resistance did not explain the effect size of land management type on λ (Appendix S4, Figure S2). However, flowering duration was positively related to the effect size of land management type on λ; species with lower flowering duration (i.e. *Plantago lanceolata, Bromus erectus, Tragopogon orientalis*) performed better in the mowing treatment while those with longer flowering duration (*Dianthus carthusianorum, Scabiosa ochroleuca*) performed better in the grazing treatment (t = 6.087, p < 0.001, r^2^ = 0.8; Figure. 3a). The start of flowering was a poor predictor of management preferences (Figure 3b, t = −0.206, p = 0.842, r^2^ = −0.1191). of flowering for each species. The line indicates the trend from the linear model. Symbols show the management treatment in which phenology was measured.

**Figure 3.**
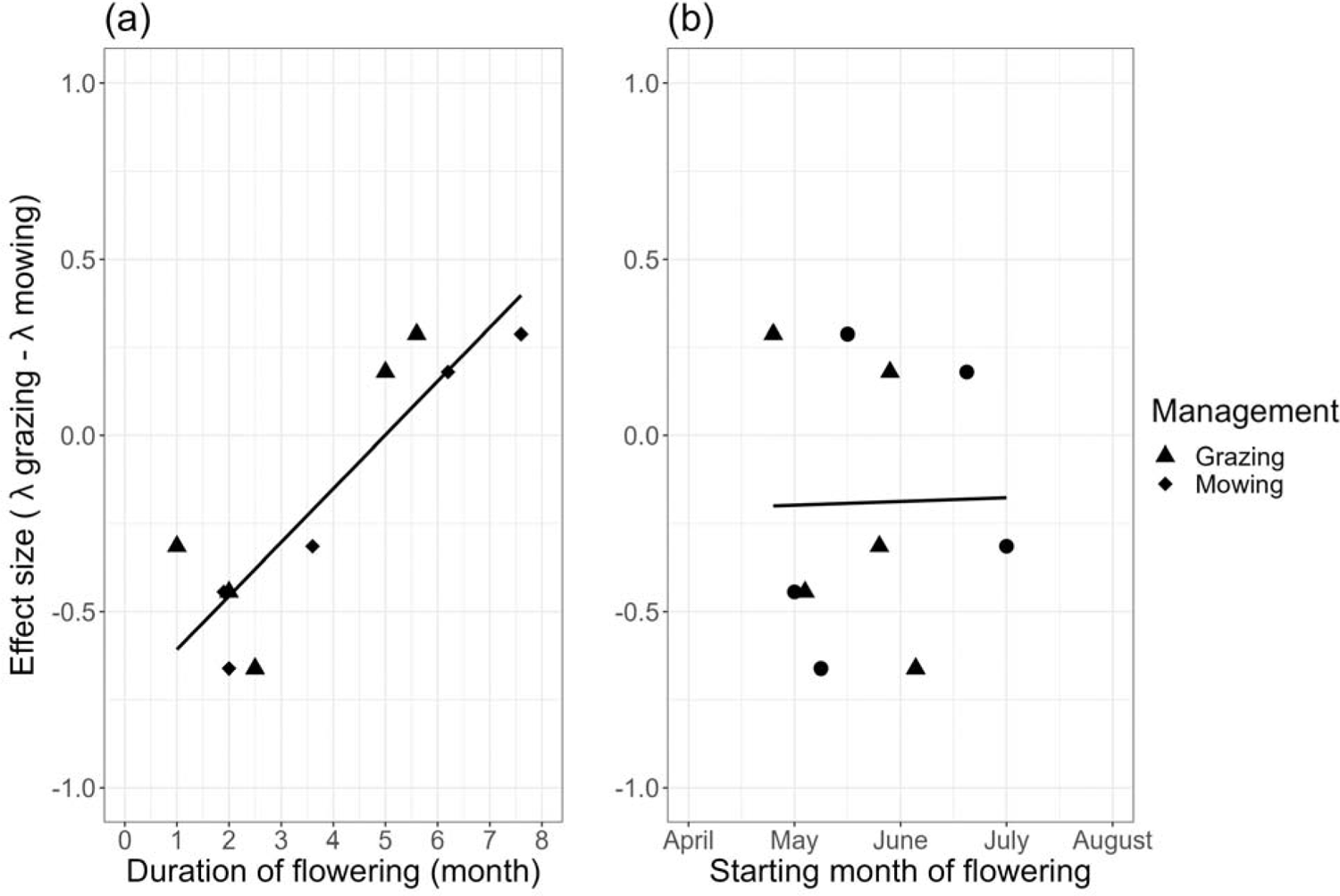
Relationship between the effect size of management on population growth rate (λ grazing – λ mowing) plotted against a) The mean duration of flowering for each species and b) the mean beginning

### Climate effects on population growth rate

*B. erectus* had a significantly higher λ in future compared to ambient climate (permutation test, p = 0.023, Figure 4a), and the other four species did not show significant differences in λ with regard to the climate treatment (Figure 4b-e).

**Figure 4.**
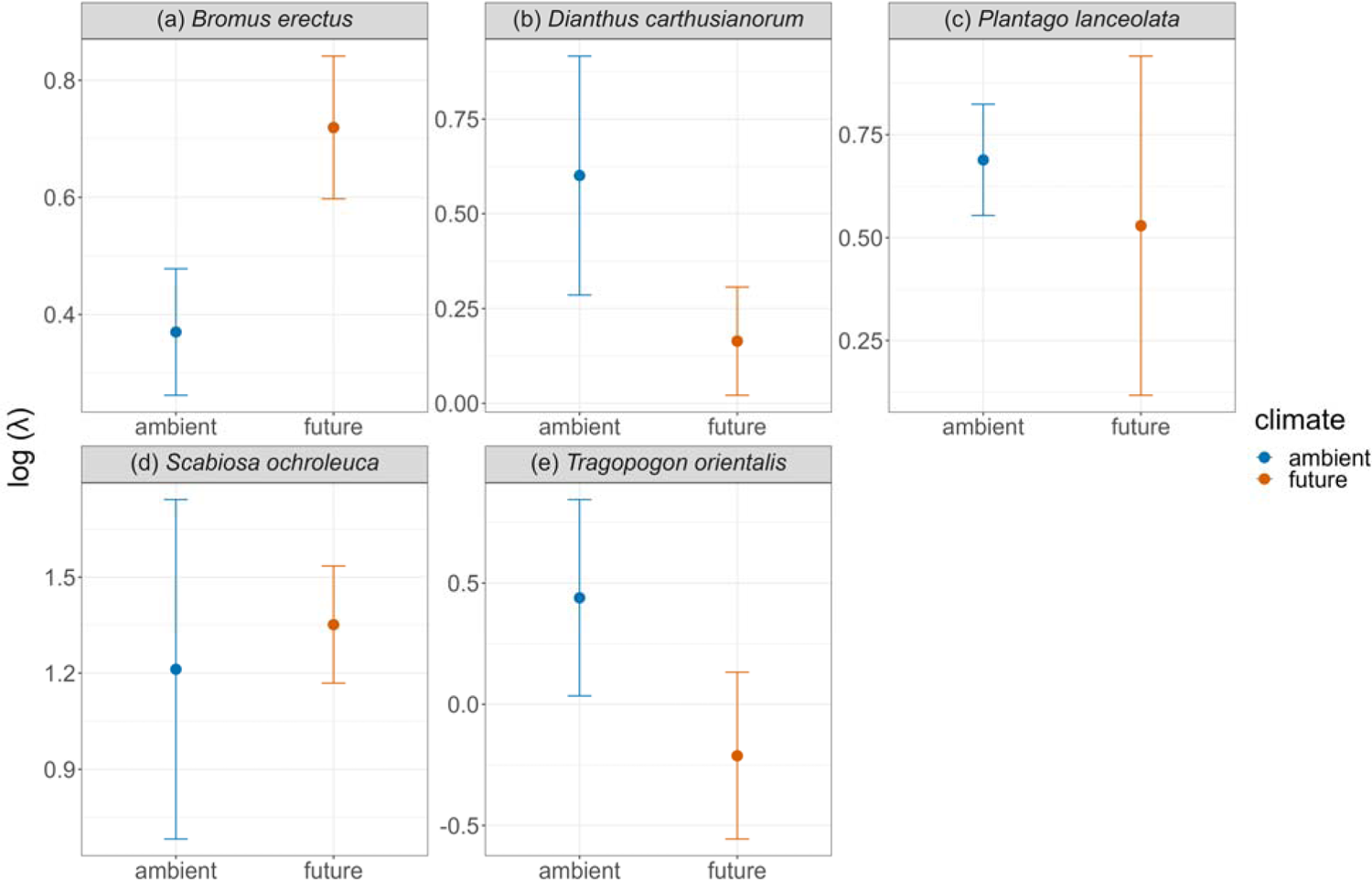
Bootstrapped logged J plotted against the two climate treatments (ambient and future) for the five different study species (a-e). Displayed is the mean logged J of the 1000 repetitions, with the standard deviation.

### Effect of climate and land management type on population growth rate

*B. erectus* had a significantly higher λ in the ambient mowing than in the ambient grazing treatment combination (permutation test, p = 0.001). In *S. ochroleuca,* λ was significantly higher in future compared to ambient climate in the mowing land management type (future mowing > ambient mowing; permutation test, p = 0.011) but not in grazing land management (permutation test, p > 0.05, Figure 5d). The λ of *T. orientalis* was significantly higher in ambient compared to future climate only in the grazing management treatment (ambient grazing > future grazing; permutation test, p = 0.002) and λ was higher in mowing compared to grazing management in both climate treatments (permutation test, ambient grazing > ambient mowing, p = 0.048, future grazing > future mowing, 0.023 Figure 5e). For all other species (*B. erectus, D. carthusianorum* and *P. lanceolata*) the lines in the interaction plots crossed but they were not significantly different from each other due to the large variability around the mean (Figure 5a-c).

**Figure 5.**
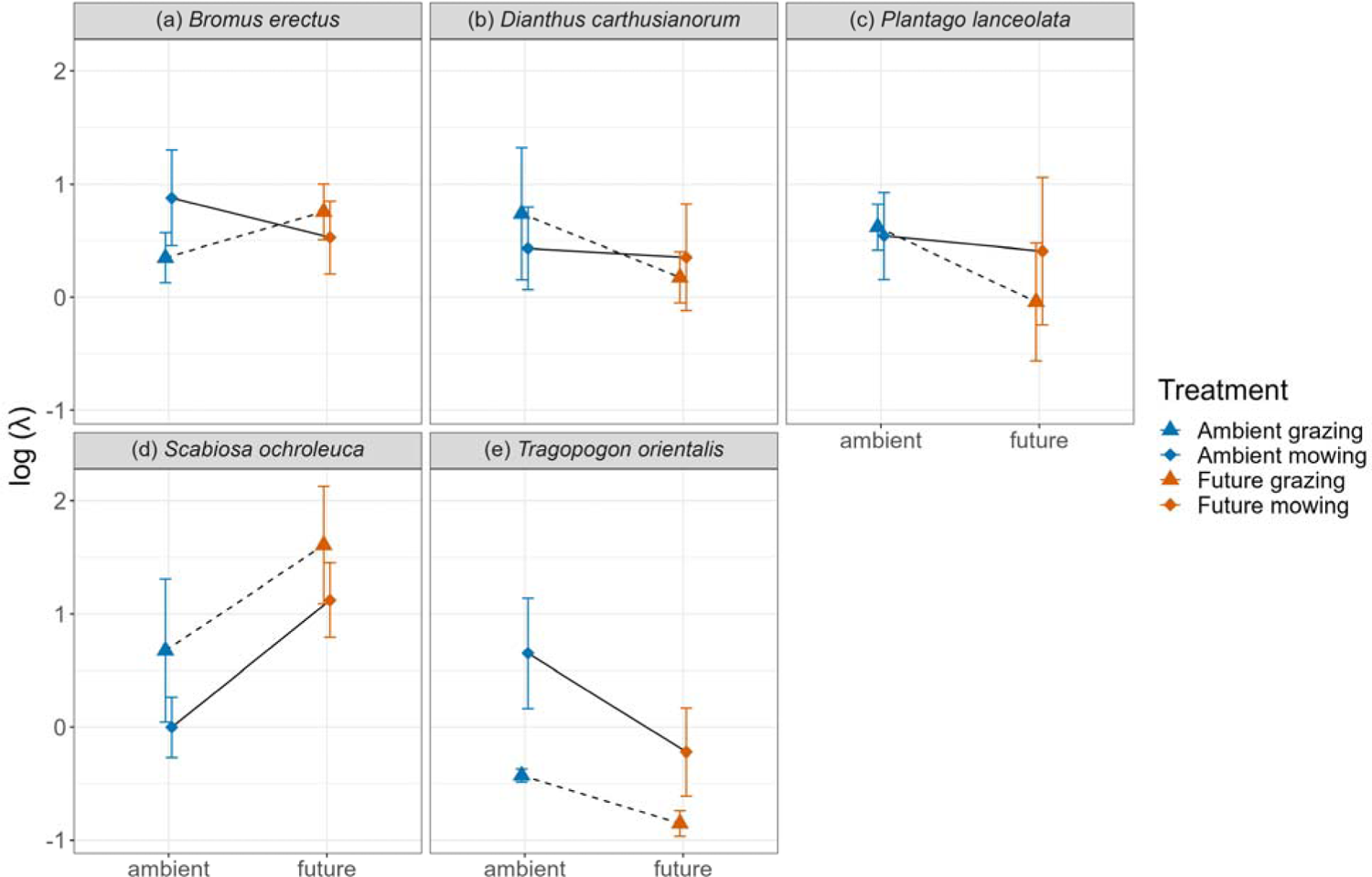
Effect of climate (ambient and future) and land management type (grazing and mowing) on population growth rate (J) of the five study species (a-e). Displayed is the mean and the standard deviation.

### Sensitivity

Across all species and treatments, λ was most sensitive to changes in reproduction parameters (Figure 6). While the magnitude differed between treatments and species (e.g. *S. ochroleuca* highest in grazing and future *D. carthusianorum* highest in ambient).

**Figure 6.**
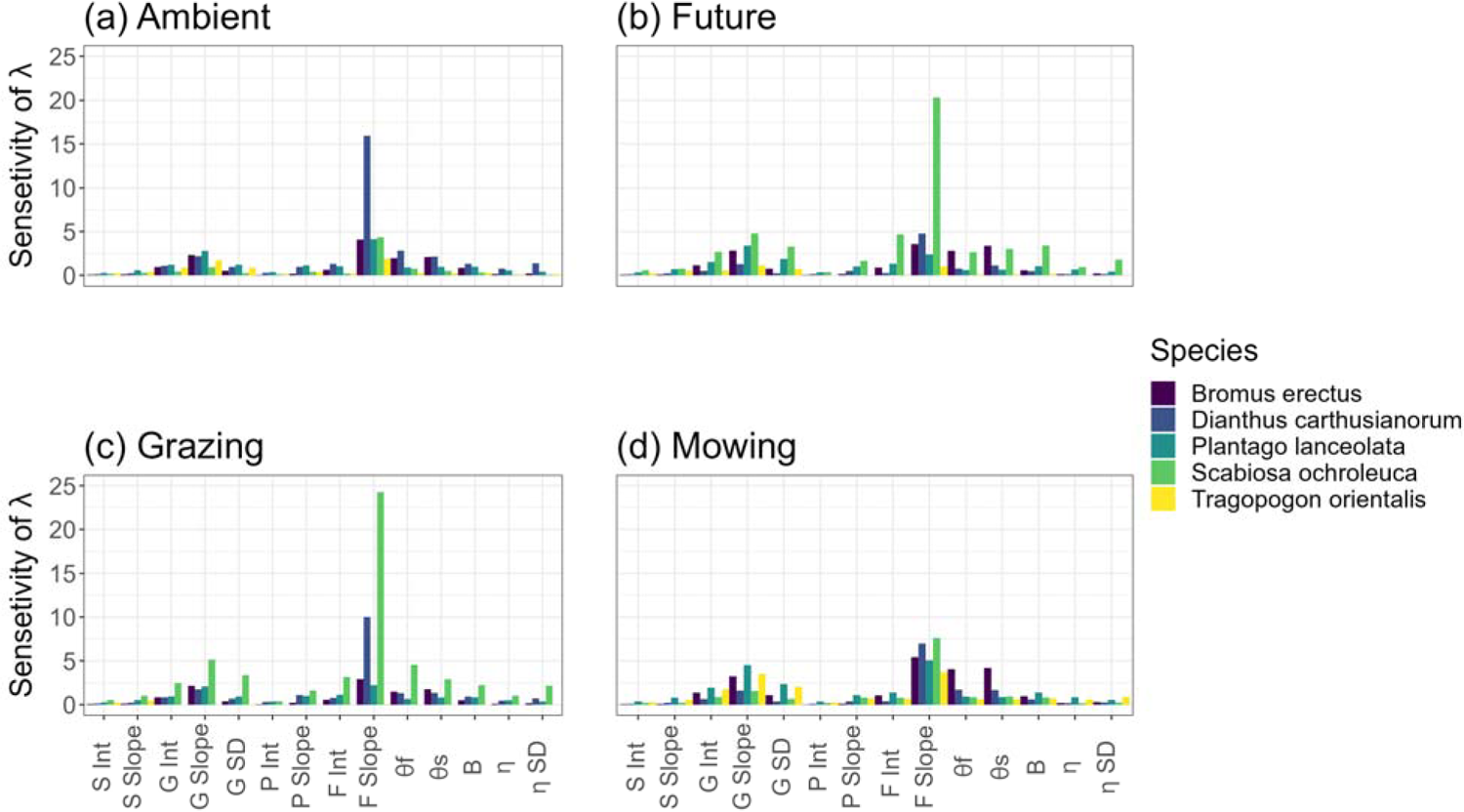
Sensitivity of pairwise treatment combinations for every vital rate included in the IPM. Each species is color-coded.

### Life Table Response Experiment

The higher λ of *B. erectus* in future compared to ambient climate was primarily due to increases in reproduction and recruitment of this species under future climate (Figure 7c and 7d). The higher λ of *S. ochroleuca* in future compared to ambient climate in the mowing management treatment was primarily due to increases in the reproduction of this species in the future mowing treatment (Figure 7d). The higher λ of *T. orientalis* in ambient compared to future climate in the grazing management treatment was primarily due to increases in plant survivorship (Figure 7c), whereas the higher λ in this species in mowing compared to grazing in future climate treatment was due to higher survival and reproduction (Figure 7b).

**Figure 7.**
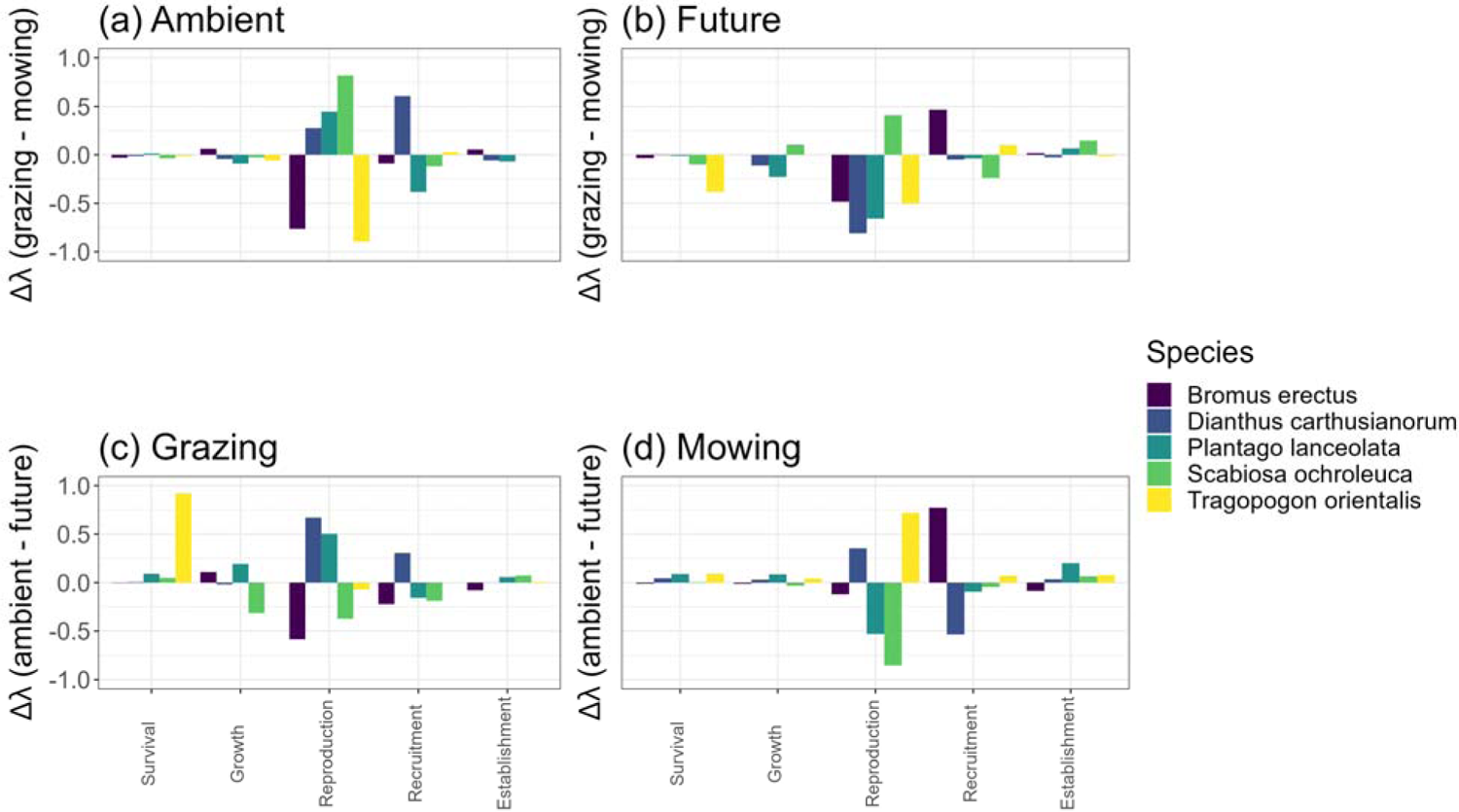
Results for the Life Table Response Experiment (LTRE). Each species is displayed in a different color. a) Ambient climate treatment and the effect of the two land management types on the parameters. b) Future climate treatment and the effect of the two land management types on the parameters. c) Grazing land management type and the effect of climate on the parameters. d) Mowing land management type and the effect of climate on the life stages. For a better overview, we summed the parameters according to their corresponding live stage.

## Discussion

Our study aimed to examine whether plant species interactively respond to climate and land management types in their demography and population dynamics. However, in the early years of our study, six of our eleven focal species went quasi extinct from the experiment, likely due to the extreme drought conditions that occurred in our region in 2018 and 2019. This is becoming an increasing phenomenon in experimental global change research—even the control treatments experience extreme climate events because these are now more common in our rapidly changing world (IPCC, 2014; Jentsch et al., 2007; Orlowsky & Seneviratne, 2012; Tebaldi et al., 2006). Of the remaining five species, we found that flowering duration is an important trait that predicts responses to land management types, with shorter duration species having higher λ in mowing management and longer duration species having higher λ in grazing management. In contrast to our expectation that plants would perform better in ambient compared to future climate conditions, we found that two species (*B. erectus* and *S. ochroleuca*) have higher λ in the future climate treatment and only *T. orientalis* shows the expected higher λ in ambient climate. We suggest this is likely due to the high drought tolerance of *B. erectus* and *S. ochroleuca*. We did not find strong support that the effects of climate and land management types were interactive for our study species, and it is unclear if this is because these interactive effects do not occur or because we have low power to detect them.

We expected that plants species in the study region (Central Germany) would perform best in ambient climate, as they would have difficulty dealing with the dry conditions expected in the summer under future climate. However, our results do not support this hypothesis. The first year of our demographic study, 2018, coincided with the driest summer for Germany since start of recording (Schuldt et al., 2020). This drought had dramatic and long-lasting effects on the plant communities across the entire GCEF experiment. Many of the species that went quasi-extinct were those that are indicative of moist soil conditions. Four of the five species that remained are indicative of drier habitat conditions and commonly occur in semi-dry or dry meadows (*B. erectus, D. carthusianorum* and *S. ochroleuca*) or are known to be highly generalized with regards to the moisture of the habitat (*P. lanceolata*). *B. erectus* and *S. ochroleuca* were also the two species that performed better in future experimental climate conditions. This result matches previous experimental studies (Lemmer et al., 2021; Moser et al., 2011; Poniatowski et al., 2018) that show high performance of *B. erectus* in hot and dry summer conditions. Our LTRE results revealed that the higher λ of *B. erectus* in future climate was due primarily to the higher reproduction and recruitment in this treatment (see also Lemmer et al. 2021). The earlier flowering of *B. erectus* compared to other species could explain its relative advantage in future climate conditions; *B. erectus* is able to take advantage of the additional water resources in spring and reproduce before the hot and dry summer conditions in the future climate treatment.

Only one of our remaining species, *T. orientalis,* is indicative of moist to moderately dry soil conditions (Sebald et al., 1990; Ellenberg et al., 1991; Bomble, 2013) and managed to persist in our study. This species did indeed show the expected higher performance in the ambient compared to future climate treatment. We suggest that we may have found more support for our hypothesis if we had been able to study the demography of more typical species from moister grassland communities as we initially planned. Our study highlights the importance of historical legacies in shaping experimental results and support projections that communities may shift towards more drought tolerant species under future climate conditions (Belovsky & Slade, 2021; Craine et al., 2013).

Surprisingly, most of our study species did not show strong differences in their population growth across land management types. The only exception was the biennial species, *T. orientalis*, which performed significantly better in the mowing compared to grazing management treatment, primarily due to higher survival and reproduction in the mowing treatment. The vegetative parts of *T. orientalis* are relatively soft and easy for sheep to chew (personal observation) and the large flower heads may attract sheep. We also observed that the flower buds were eaten by other wild animals, such as rabbits (personal observation). This species adds to several studies that showed that grazing has a negative effect on plant population dynamics (Hansen & Wilson, 2006; Jacquemyn et al., 2012; van der Meer et al., 2014). For example, in *Primula veris*, grazing was found to decrease population growth rate, primarily through its negative effect on reproduction (Jacquemyn et al., 2012).

Our results show that the effect of management on λ depended on flowering duration. Plants that flowered a relatively short duration showed higher λ in mowing (e.g. *T. orientalis*), whereas those flowering across a longer duration showed higher λ in the grazing management. This could be because plant populations that flower across a long time period are buffered from losing all of their reproductive success from disturbances that occur in a short time interval, such as grazing. However, we have to be cautious when interpreting these results as in our study years the management was mostly applied in a lower frequency than usual; most years had one mowing and two grazing events. This was because mowing and grazing were not possible when there was too little biomass regrowth, which occurred during the dry study years. Other studies have also shown that low frequency (once a year) and late season mowing benefits the reproduction of plants (Jantunen et al., 2007; Johansen et al., 2019). Across all of our study species, we found that the population growth rates were most sensitive to reproduction. Thus, it may be a general finding that management treatments should be timed to interfere as little as possible with peak flowering and fruiting (Nakahama et al., 2016; Reisch & Poschlod, 2009).

We expected that interaction effects of treatments on λ would be common in our study and that each species would have a treatment combination that was most optimal for λ. However, we had little power to test for these interactions due to the high variation across individuals in the sensitive vital rates of seed production and recruitment. For example, the high variation in λ of *S. ochroleuca* in the ambient-grazing treatment combination is likely due to very high seed production (>2000 seeds) of three individuals in this treatment combination (Appendix S6). Similarly, the high variation in λ in the ambient treatment for *D. carthusianorum* is likely due to high seed production of a few individuals in the ambient mowing treatment (Appendix S6) and high recruitment of new spring seedlings in a few plots in the ambient grazing treatment (Appendix S7). Finally, while *P. lanceolata* appears overall have higher λ in ambient compared to future climate treatments, high variance in the future-mowing treatment combination, which is likely due to two reproductive individuals with outlier high seed production, may prevent the detection of an effect. However, it is noteworthy that across all species in this study many of the plants with outlier seed production were in the ambient climate treatment, suggesting that the future climate treatment might negate the positive effects that a preferred management treatment might have on the reproduction and population growth rate of these grassland species.

In demographic studies of both plants and animals, reproductive output is known to be skewed across individuals, with only a few individuals producing many offspring (van Daalen & Caswell, 2020). The ability to distinguish deterministic treatments effects from stochasticity among individuals would require much higher sample sizes than is available in our GCEF experiment. While there are many advantages of conducting multi-species demographic studies like this one and of conducting demographic research in a controlled experimental setting, one distinct disadvantage is the effort and sample size required to adequately detect small changes in sensitive vital rates (McMahon et al., 2019).

It was surprising to find in our LTRE that changes in reproduction and recruitment often contributed most to the observed responses of λ to management and climate. Usually in longer lived species, λ is highly sensitive to changes in survival (Weppler, Stoll and Stöcklin, 2006; Kuss et al., 2008) and thus even small changes in survival across treatments should contribute significantly to changes in λ. In our case, we find that λ is highly sensitive to recruitment. This might be due to the relatively high λ of our perennial species; species with high λ are often more sensitive to reproduction and recruitment than to survivorship (Franco & Silvertown, 2004; Ramula et al., 2008). Further, many of our species were able to maintain their survival across treatments, but had high changes in their reproduction and recruitment.

## Conclusions

Our study is a first step towards answering important yet unanswered question on how climate change affects population dynamics of grassland species and how this effect is modified by different land management types (Ehrlen, 2019). Our results are optimistic for our four more drought-tolerant species characteristic for dry or semi-dry grasslands, showing persistent or growing populations in both management types (grazing vs. mowing) and under future climate conditions as well as after natural extreme drought events. However, less drought-tolerant species either went quasi extinct early in the study or were projected to decline in the future, indicating that populations cannot persist in the presence of future climate conditions. Our results suggest that it will be more difficult to create and maintain especially mesophilic grasslands composed of species with preference for moister soil conditions in the future. Furthermore, we suggest that management in future climate should carefully be timed outside of peak flowering and fruiting, so that as many species as possible can reproduce and persist in managed grasslands.

## Supporting information

Full Appendix

## Acknowledgements

This research was funded by the Alexander von Humboldt Foundation (Alexander von Humboldt Professorship of TMK) and the Helmholtz Recruitment Initiative of the Helmholtz Association (TMK). The authors gratefully acknowledge the support of iDiv funded by the German Research Foundation (DFG-FZT 118). We appreciate the Helmholtz Association, the Federal Ministry of Education and Research, the State Ministry of Science and Economy of Saxony-Anhalt, and the State Ministry for Higher Education, Research and the Arts Saxony, which fund the Global Change Experimental Facility (GCEF) project. We thank the staff of the Bad Lauchstädt Experimental Research Station (especially Ines Merbach and Konrad Kirsch), for their work in in maintaining the plots and infrastructures of the GCEF, and François Buscot, Stefan Klotz, Thomas Reitz, and Martin Schädler for their role in setting up the GCEF. We thank Julia Lemmer, Christian Savona, Simon Bitzan, Georg Küstner, Lina Lüttgert and Amibeth Thompson for their help with the fieldwork.

## Author contributions

The study was designed by LK (leading) and MA and TMK (supporting). Fieldwork was designed and carried out by LK and MA (both leading). The data for the phenology of plants was collected by CP (leading). Analysis was carried out by MA (leading) with support of the other co-authors. The manuscript was written by MA (leading) and TMK and LK (supporting). All authors commented and improved the manuscript.

