## Supplementary material for "Reproductive success mediates the effects of climate change and grassland management on plant populations dynamics": Full Appendix

**Ecology**

**Appendix S1**

**Figure S1** Layout of every GCEF plot. 1 indicates where the transect was situated at each GCEF subunit. Displayed is the overall size of the GCEF subunits and the inner subunit where experiments take place.

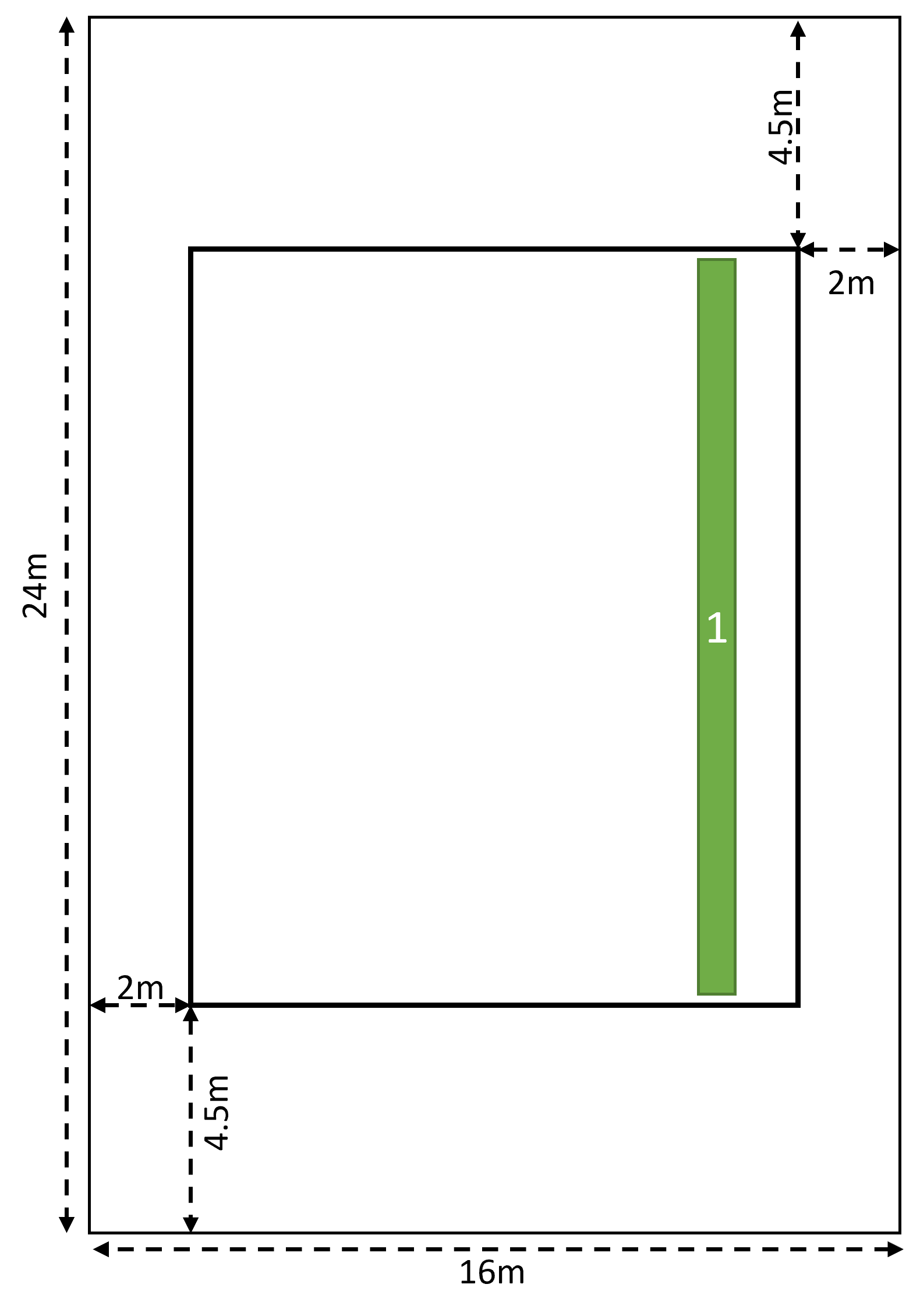

**Appendix S2**

**Table S1** Given are the dates of the management events. Sheep graze one plot for 24 hours and thus the management took place over the course of 10 days. The sheep were let on one plot in the morning of the day and the next day they moved to the next plot. Mowing was done in one day.

| **Management event** | **Year** | **Date** |
| --- | --- | --- |
| Grazing | 2018 | 30.04 - 11.05 |
| Grazing | 2018 | 11.06 - 22.06 |
| Mowing | 2018 | 11.06 |
| Grazing | 2019 | 6.05 - 17.05 |
| Grazing | 2019 | 10.06 - 21.06 |
| Mowing | 2019 | 11.06 |
| Grazing | 2020 | 11.05 - 22.5 |
| Grazing | 2020 | 22.06 - 03.07 |
| Mowing | 2020 | 08.06 |
| Grazing | 2021 | 10.05 - 21.05 |
| Grazing | 2021 | 14.06 - 25.06 |
| Mowing | 2021 | 14.06 |
| Mowing | 2021 | 06.09.2023 |
| Grazing | 2021 | 06.09 - 17.09 |

**Appendix S3**

**Table S2** Information on how we identified seedlings. Mostly that stage class was size dependent and if an individual plant didn’t grow to a certain size, it was identified as seedling. Some Bromus erectus individuals flowered after they developed 4 leaves but still were smaller than 0.25 cm^2^ which is why we classified every individual with three or more leaves as adult.

| **Species** | **Maximum size as a seedling** |
| --- | --- |
| Anthoxanthum odoratum | < 4 leafs |
| *Bromus* *erectus* | 1 ramet with maximum 3 leafs |
| Crepis bunnies | 1 ramet with maximum 3 leafs |
| *Dianthus* *carthusianorum* | Size of rosette <0.25cm^2 |
| Lotus corniculatus | Size of rosette <0.25cm^2 |
| Lychnis flos-cuculi | Didn’t find seedling |
| Medicago falcata | Size of rosette <0.25cm^2 |
| *Plantago* *lanceolata* | 1 ramet with maximum 3 leafs |
| *Scabiosa* *ochroleuca* | 1 ramet with maximum 2 leafs |
| *Tragopogon* *orientalis* | Maximal 3 leafs |
| Trifolium pratense | 1 individual with maximum 2 leafs |

**Appendix S4**

**Figure S2** Grazing and mowing resistance plotted against the effect size of $\lambda$. Negative values indicate that the species has a higher $\lambda$ in the mowing treatment. Panel a.) shows the effect of grazing resistance on the effect size and panel b.) shows the effect of mowing resistance on the effect size.

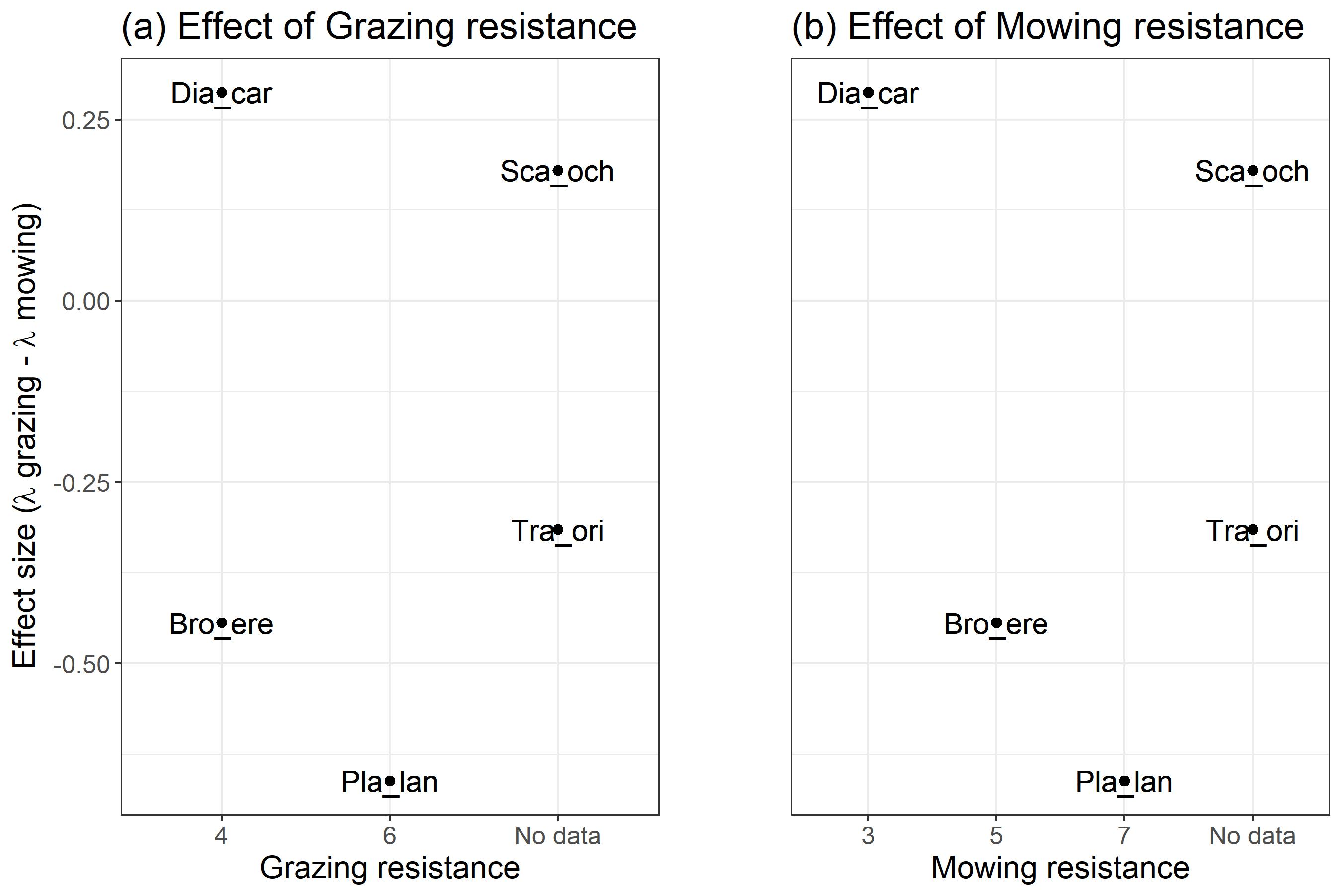

**Appendix S5**

**Figure S3** Bootstrapped λ of each species over the study years for the management treatment (mowing, grazing). Displayed is the mean λ and the standard deviation. Each year on the x axis stands for a transition: 2018 = 2018–2019, 2019 = 2019–2020, 2020 = 2020–2021, 2021 = 2021–2022. For the transition 2021 we were not able to calculate a lambda for T. orientalis in the grazing land management treatment due to lack of data points.

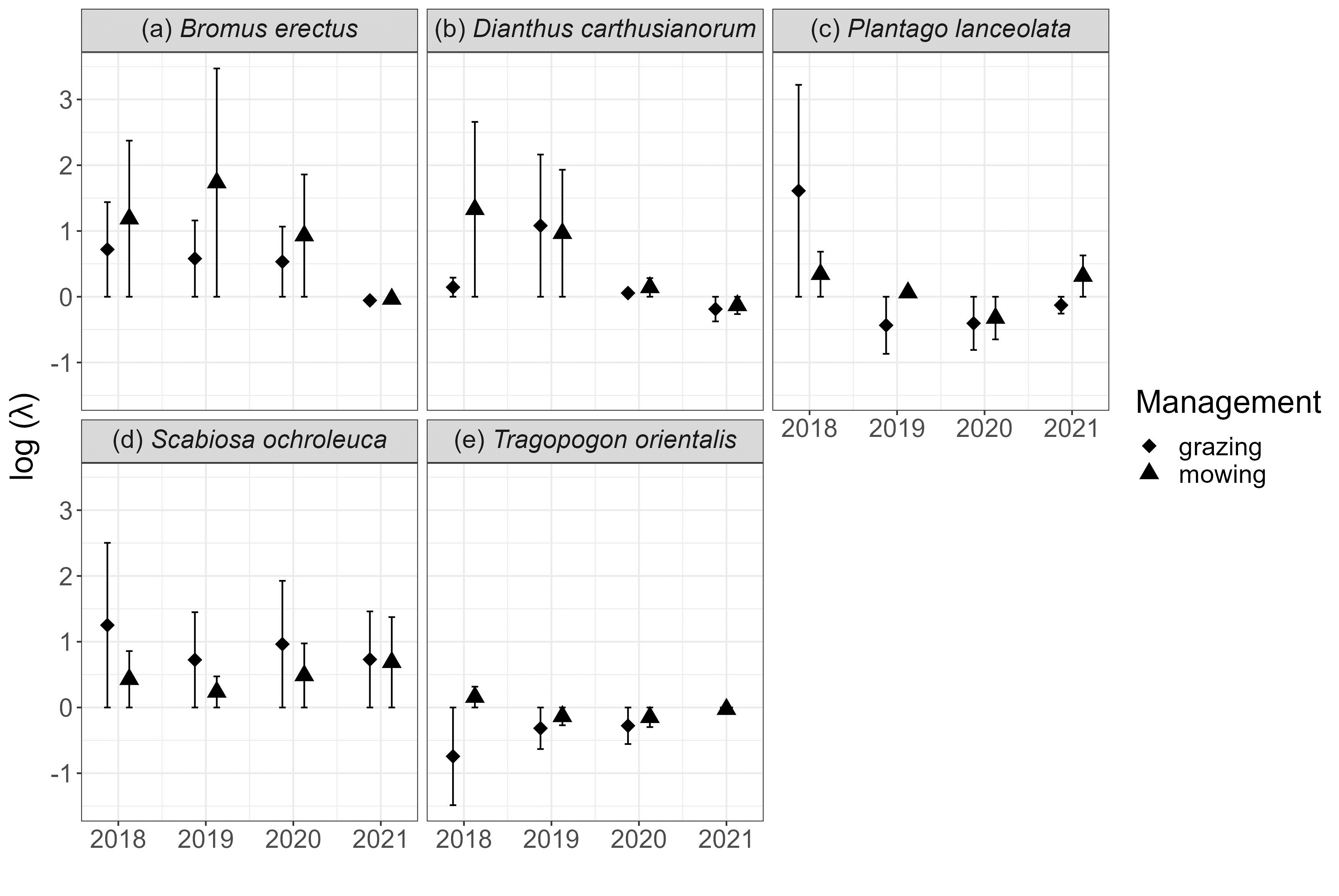

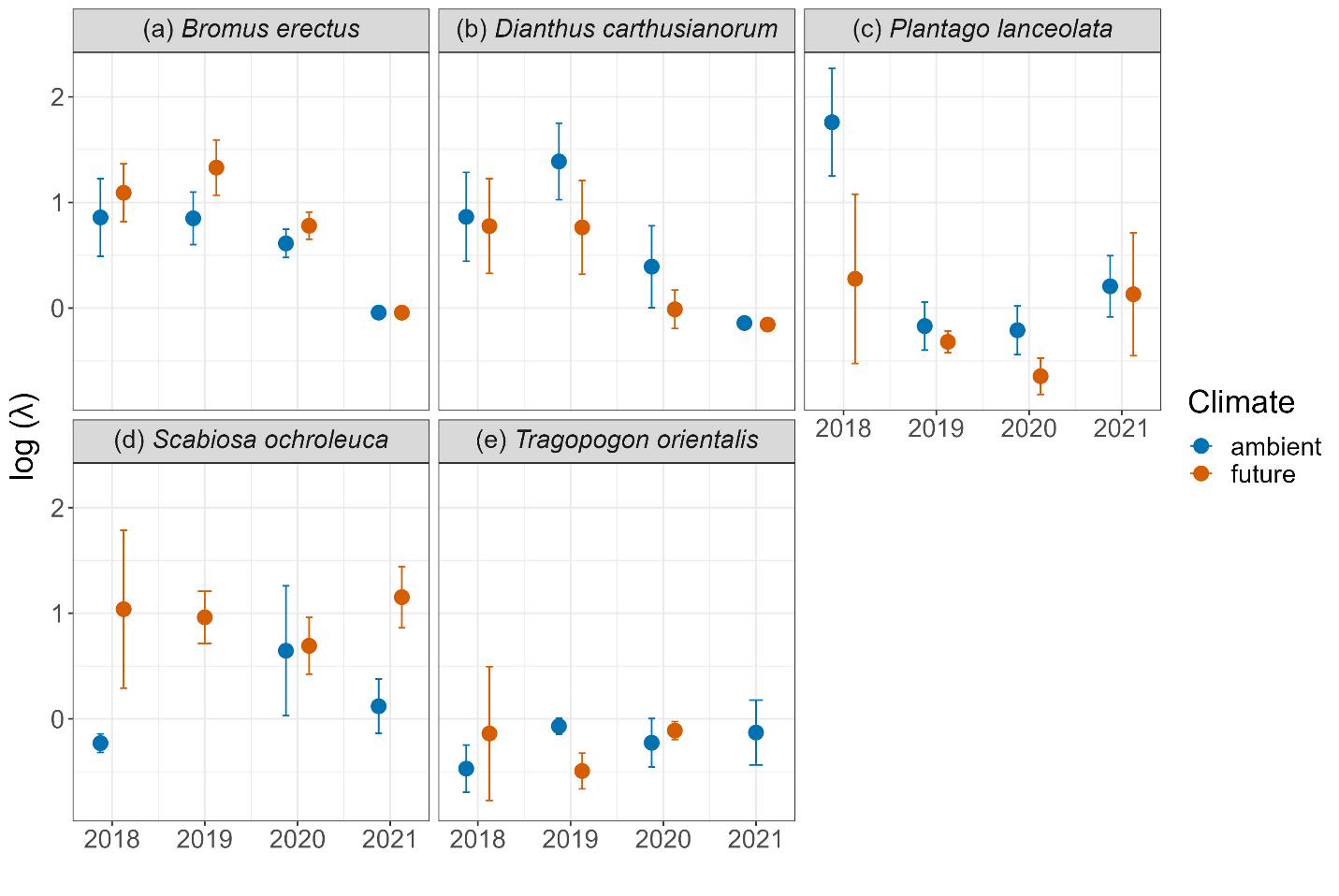
**Figure S4** Bootstrapped λ of each species over the study years for the climate treatment (ambient, future). Displayed is the mean λ and the standard deviation. Each year on the x axis stands for a transition: 2018 = 2018 – 2019, 2019 = 2019 – 2020, 2020 = 2020 – 2021, 2021 = 2021 – 2022. *S. ochroleuca* in 2019 and *T. orientalis* did not have enough data points in order for us to calculate the population growth rate.

**Figure S5** The effect of the treatment combinations (management and climate) over the years of the study on λ. Displayed is the mean λ and the standard deviation. Each year on the x axis stands for a transition: 2018 = 2018 – 2019, 2019 = 2019 – 2020, 2020 = 2020 – 2021, 2021 = 2021 – 2022. Missing points are due to lack of data points to calculate the population growth rate.

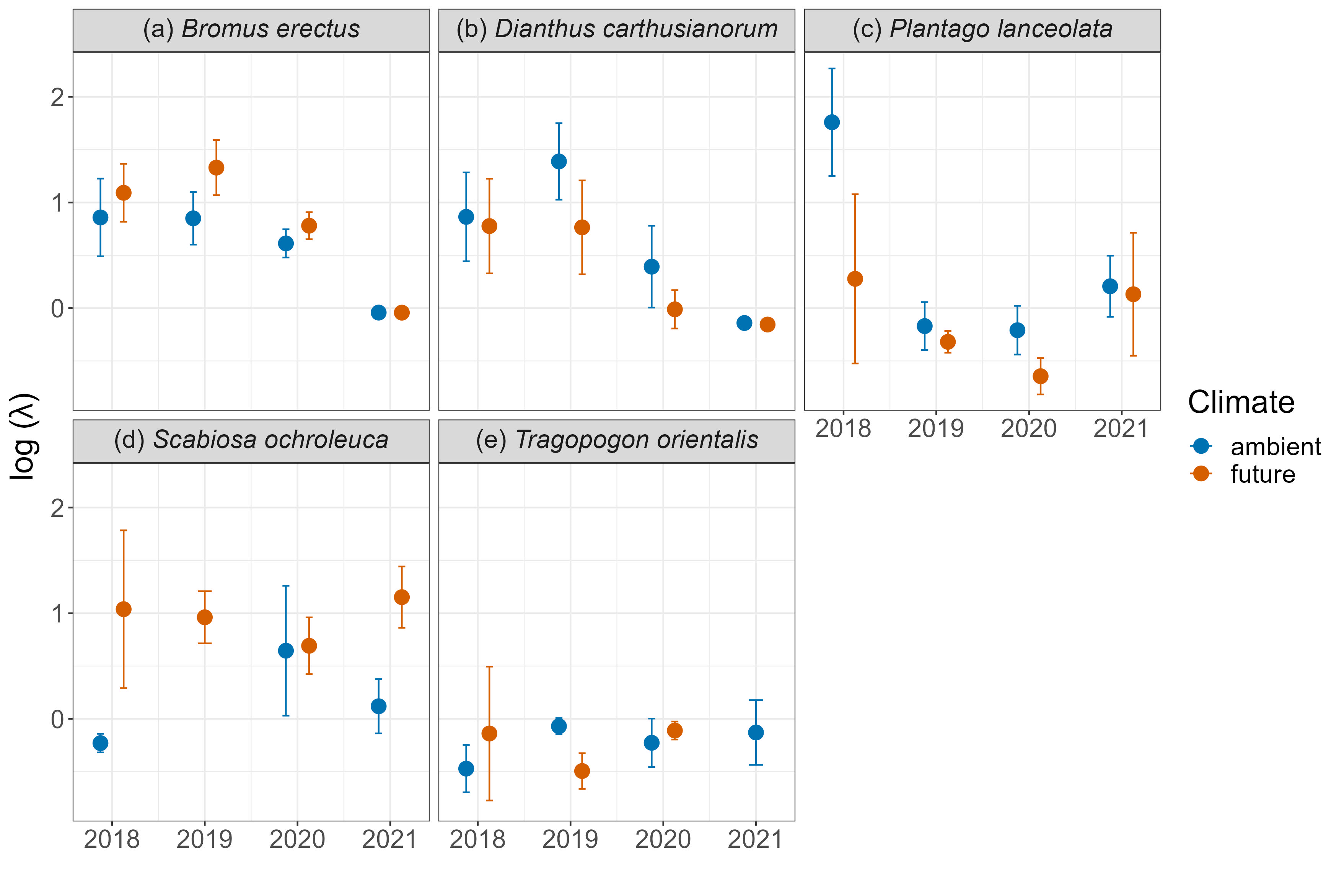

**Appendix S6**

**Figure S6** Number of seeds plotted against the size of each reproductive individual for the species (A) Bromus erectus, (B) Dianthus carthusianorum, (C) Plantago lanceolata, (D) Scabiosa ochroleuca and (E) Tragopogon orientalis. Different treatments are indicated with different shapes and colors.

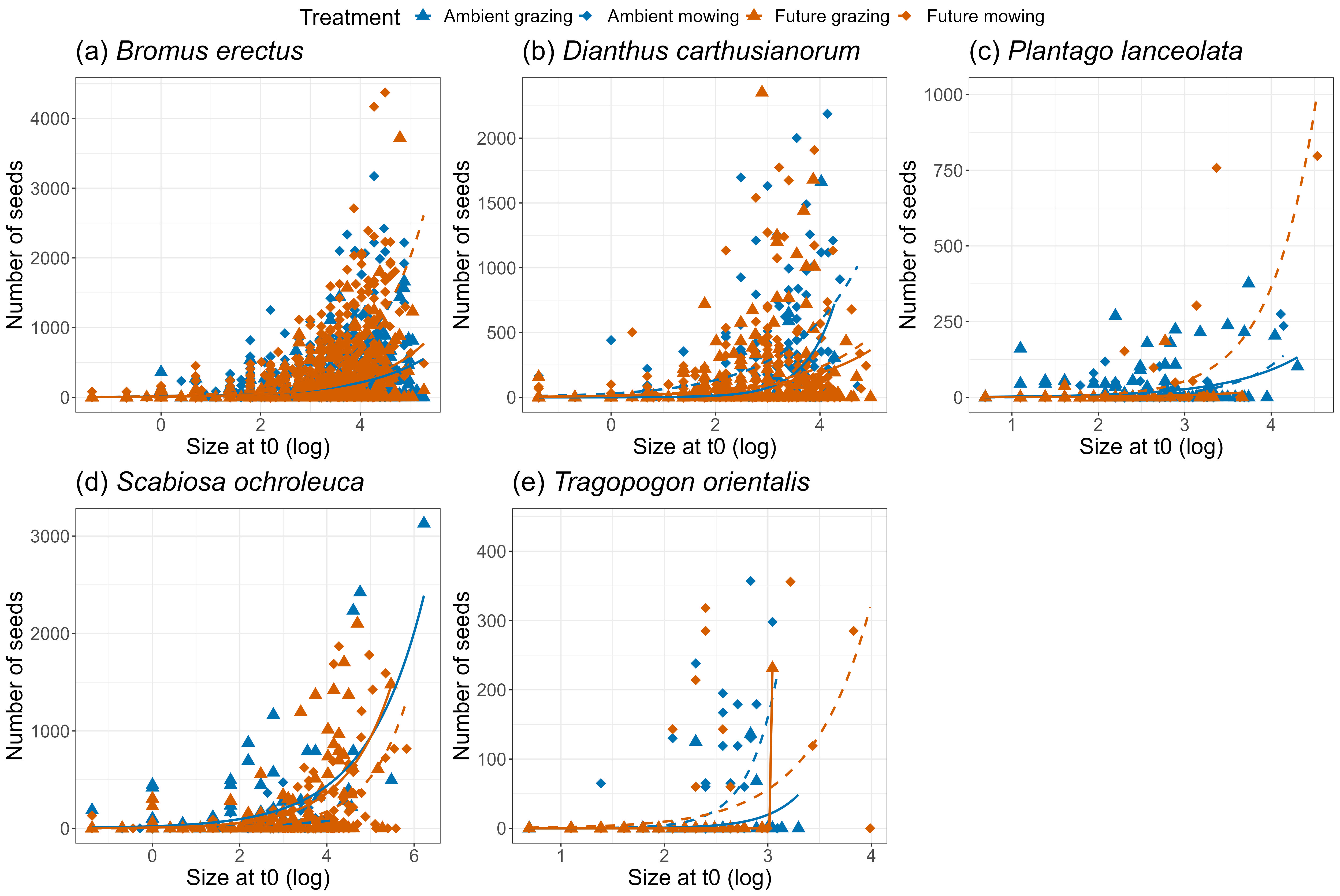

**Appendix S7**

**Figure S7** Turnover from seeds to seedling in the different treatments in spring (spring recruitment) for (a) Bromus erectus (b) Dianthus carthusianorum (c) Plantago lanceolata (d) Scabiosa ochroleuca

(e) Tragopogon orientalis

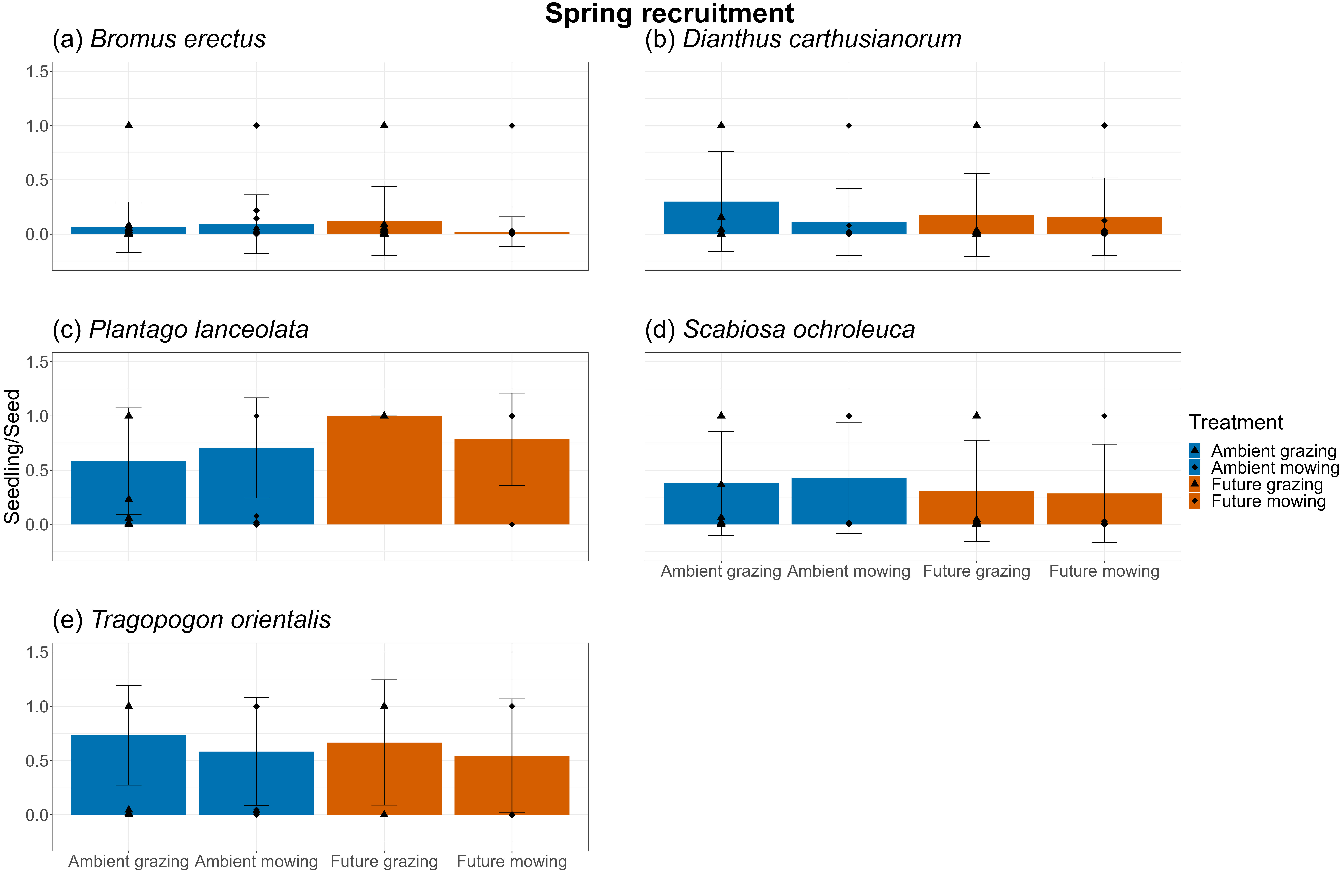

**Figure S8** Turnover from seeds to seedling in the different treatments in fall (fall recruitment) for (a) Bromus erectus (b) Dianthus carthusianorum (c) Plantago lanceolata (d) Scabiosa ochroleuca

(e) Tragopogon orientalis

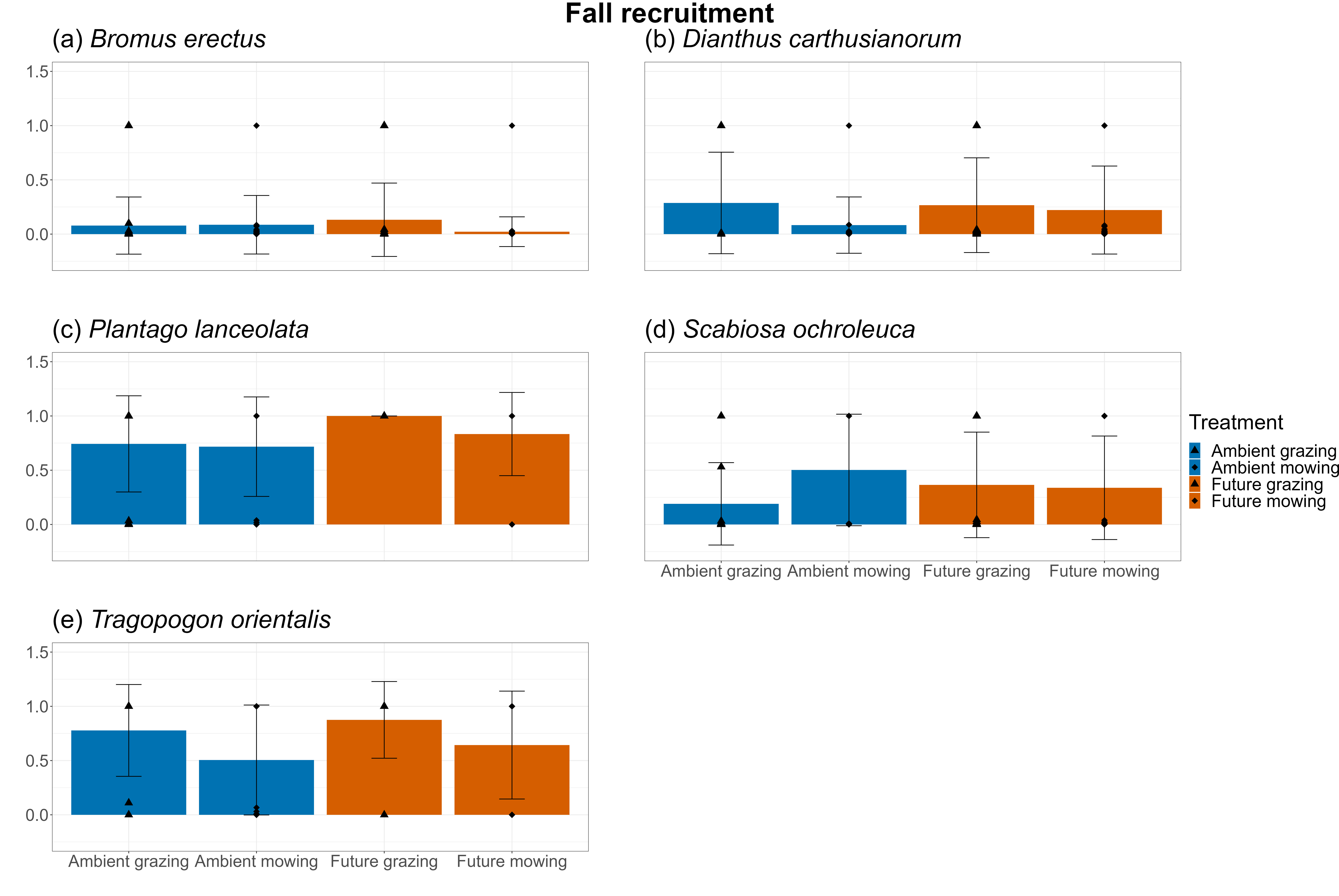
